# The Single Cell Transcriptomic Landscape of Early Human Diabetic Nephropathy

**DOI:** 10.1101/645424

**Authors:** Parker C. Wilson, Haojia Wu, Yuhei Kirita, Kohei Uchimura, Helmut G. Rennke, Paul A. Welling, Sushrut S. Waikar, Benjamin D. Humphreys

## Abstract

Diabetic nephropathy is characterized by damage to both the glomerulus and tubulointerstitium, but relatively little is known about accompanying cell-specific changes in gene expression. We performed unbiased single nucleus RNA sequencing (snRNAseq) on cryopreserved human diabetic kidney samples to generate 23,980 single nucleus transcriptomes from three control and three early diabetic nephropathy samples. All major cell types of the kidney were represented in the final dataset. Side by side comparison demonstrated cell-type-specific changes in gene expression that are important for ion transport, angiogenesis, and immune cell activation. In particular, we show that the diabetic loop of Henle, late distal convoluted tubule, and principal cells all adopt a gene expression signature consistent with increased potassium secretion, including alterations in Na-K^+^-ATPase, *WNK1*, mineralocorticoid receptor and *NEDD4L* expression, as well as decreased paracellular calcium and magnesium reabsorption. We also identify strong angiogenic signatures in glomerular cell types, proximal convoluted tubule, distal convoluted tubule and principal cells. Taken together, these results suggest that increased potassium secretion and angiogenic signaling represent early kidney responses in human diabetic nephropathy.

**Significance Statement:** Single nucleus RNA sequencing revealed gene expression changes in early diabetic nephropathy that promote urinary potassium secretion and decreased calcium and magnesium reabsorption. Multiple cell types exhibited angiogenic signatures, which may represent early signs of aberrant angiogenesis. These alterations may help to identify biomarkers for disease progression or signaling pathways amenable to early intervention.

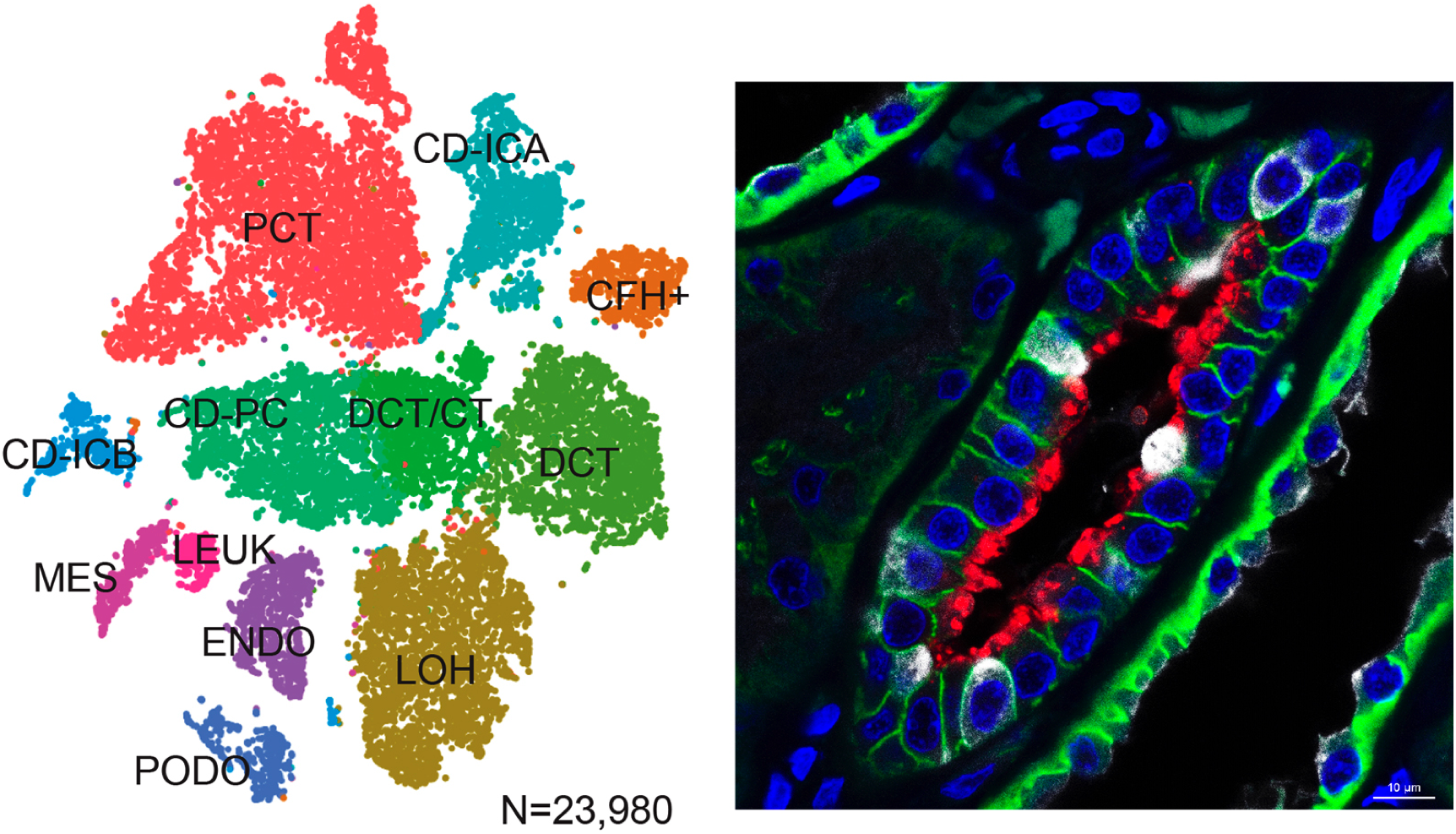

## Introduction

Single cell RNA sequencing (scRNA-seq) enables quantification of gene expression in individual cells (1). An advantage of scRNAseq over bulk RNA-seq is that it can interrogate multiple cell types simultaneously while measuring changes in transcriptional states and signaling pathways in a cell-type-specific manner. Advances in library preparation and isolation techniques, like single nucleus RNA-sequencing (snRNAseq), have increased sensitivity such that rare cell types and states can be captured from cryopreserved samples (2). We hypothesized that snRNAseq of kidney cortex in early diabetic nephropathy would reveal altered signaling pathways and gene expression patterns that would reflect the earliest adaptive changes to hyperglycemia.

Although diabetic nephropathy is the leading cause of end-stage renal disease, relatively little is known about early transcriptional changes that precede overt diabetic nephropathy. Laboratory measures like serum creatinine and urine protein are not sufficiently sensitive to detect the earliest manifestations of diabetes, and patients may go for years before they accrue sufficient damage to come to clinical attention. Although efforts are underway to develop biomarkers for the detection of early diabetic kidney disease, there is no consensus on the best approach to stratify patients for risk of progression (3).

Early histologic signs of diabetic nephropathy include thickening of the glomerular basement membrane, mesangial expansion, and podocyte loss, which often progresses to glomerulosclerosis. However, the cell types and signaling pathways that contribute to disease progression are poorly understood (4). Previous efforts to characterize transcriptional changes in human diabetic glomeruli by bulk RNA-seq have identified important pathways, but are limited because they can only measure the integrated and averaged gene expression of multiple cell types (5–7).

Here, we describe the first snRNA-seq analysis of early human diabetic nephropathy. We identified all major cell types in the kidney cortex and infiltrating immune cells in diabetic patients. The endothelium, mesangium, proximal convoluted tubule, and late distal convoluted tubule all had an angiogenic expression signature. We also observed changes in expression of the Na+/K+-ATPase and other transport-related genes in the loop of Henle, distal convoluted tubule, and principal cells, indicative of enhanced urinary potassium secretion. These changes were accompanied by decreased expression of negative regulators of potassium secretion, *WNK1* and *NEDD4L*, and increased expression of the calcium sensing receptor, *CASR*, and its downstream effector, *CLDN16*, suggestive of decreased paracellular calcium and magnesium reabsorption. Infiltrating immune cells were markedly increased in diabetic samples and expressed recently identified markers that predict diabetic nephropathy progression. These gene expression changes may be useful for identifying early biomarkers and signaling pathways in diabetic nephropathy.

## Results

Human kidney cortex was sampled from three non-diabetic controls and three diabetics following nephrectomy for renal mass. Patient age ranged from 52 to 74 years (mean = 60 +/− 7.8y) and did not differ between groups (Table S1). Diabetics had elevated A1c (mean = 7.9 +/− 1.5%) and two of three patients had proteinuria. Baseline serum creatinine (mean = 1.06 +/− 0.23 mg/dl) was not different between groups. Two diabetic patients had an increased proportion of global glomerulosclerosis that ranged from mild (11-25%) to moderate (26-50%), whereas controls had no glomerulosclerosis (<10%). The same two diabetic patients had mild (11-25%) interstitial fibrosis and tubular atrophy (IFTA) compared to no IFTA in controls (1-10%). Vascular injury was similar between groups and ranged from mild to moderate. All diabetic patients had evidence of mesangial sclerosis and glomerular basement membrane thickening (Figure S1).

### snRNA-seq Identifies All Major Cell Types in the Kidney Cortex

A total of 23,980 nuclei passed quality control filters (Table S2: mean = 3996 +/− 1195 nuclei per sample) and had an average of 2541 genes and 6894 unique molecular identifiers per nucleus. Eleven kidney cell types and 4 immune cell types (Figure 1A) were identified by unsupervised clustering (Figure S2).

**Figure 1.**
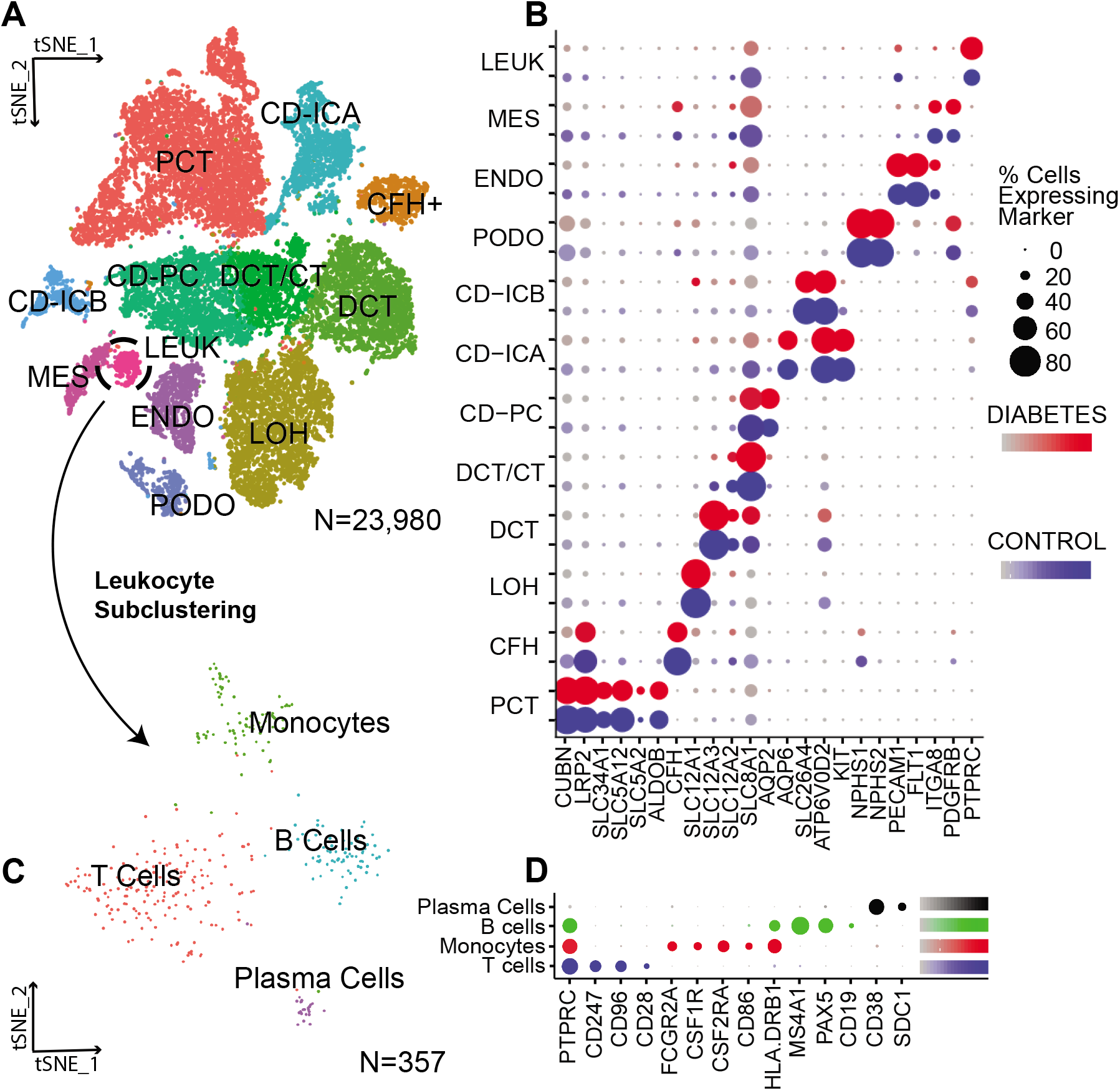
Integrated snRNA-seq dataset of diabetic and control samples. A) Diabetic and control samples were integrated into a single dataset and clustered using Seurat. PCT-proximal convoluted tubule, CFH-complement factor H, LOH-loop of Henle, DCT-distal convoluted tubule, CT-connecting tubule, CD-collecting duct, PC-principal cell, IC-intercalated cell, PODO-podocyte, ENDO-endothelium, MES-mesangial cell, LEUK-leukocyte B) Cell clusters were identified by kidney cell lineage-specific marker expression C) The leukocyte cluster (LEUK) was extracted from the integrated dataset and subclustered into leukocyte subsets D) Leukocyte subsets were identified by expression of lineage-specific markers

Clusters were annotated based on expression of lineage-specific markers (Figure 1B). We detected glomerular cell types (podocytes, mesangium, endothelium), parietal epithelium, proximal tubule, loop of Henle, early distal convoluted tubule (DCT), late distal convoluted tubule and connecting tubule (DCT/CT), and collecting duct (principal cells, type A intercalated cells, type B intercalated cells). Diabetics had an increased number of leukocytes consisting of T-cells, B-cells, mononuclear cells, and plasma cells (Figure 1C and 1D).

### Early Gene Expression Changes in the Diabetic Glomerulus

A total of 663 podocytes were evenly distributed between groups and had 57 differentially expressed transcripts (Figure 2A) enriched for gene ontology terms including neuronal cell death, phosphate metabolism, and developmental processes. Up-regulated genes included regulators of cell survival (*UNC5D, NTNG1*) (8), cell cycle and development pathways (*DACH2, EYA2, CDK6*) (9–11), response to hyperglycemia (*PDE10A, HDAC9*) (12), and glomerular basement membrane integrity (*DLC1, DTNA, ITGB3, NDRG2, MAPT*) (13). *PLA2R1* and *THSD7A*, the targets of autoantibodies in primary membranous nephropathy, were increased in diabetic podocytes, in contrast to a prior report of later stage diabetes where *PLA2R1* was downregulated sixfold (6). Patients from the prior study had more advanced diabetic nephropathy (mean sCr = 2.83 +/− 1.55 mg/dL, mean proteinuria = 1.97 +/− 0.78 g/g) compared to our cohort.

**Figure 2.**
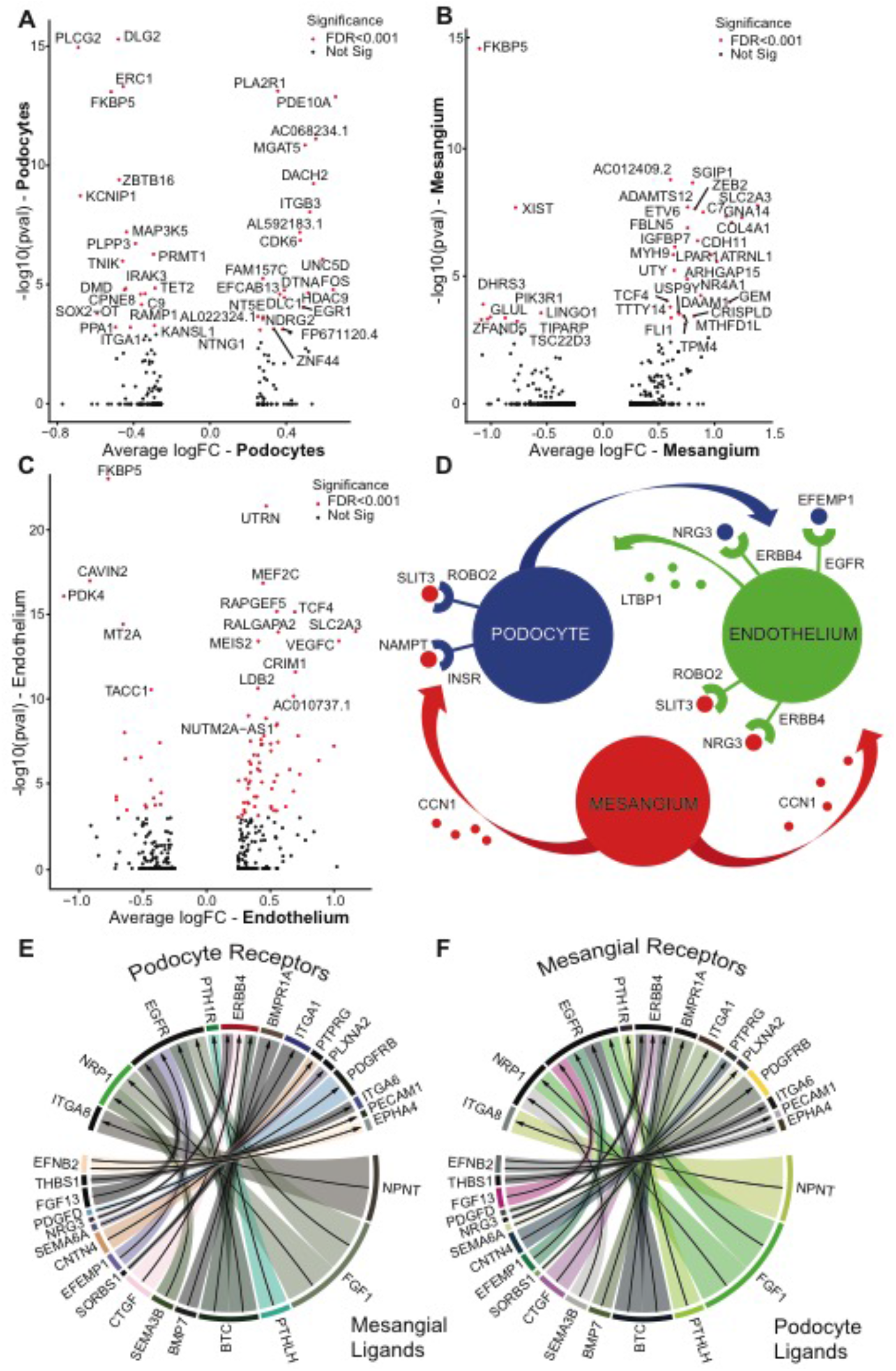
Differential gene expression and intercellular signaling in the diabetic glomerulus. - Diabetic and control samples were integrated into a single dataset and gene expression was compared within cell types using Seurat. Differentially expressed genes are displayed for A) Podocytes B) Mesangial cells and C) Endothelial cells. Ligand-receptor interactions were inferred using a publicly-available database (Ramilowski *et al*.) and include D) Differentially expressed ligand-receptor intercellular signaling pathways and E,F) All possible ligand-receptor signaling pathways in the podocyte and mesangium.

A total of 512 mesangial-like cells were identified with 45 differentially expressed genes (Figure 2B). This cluster likely represents an admixture of mesangial cells and vascular smooth muscle cells; both of which express *ITGA8* and *PDGFRB* (Human Protein Atlas). Up-regulated genes included extracellular matrix components (*COL4A1, COL4A2*), mediators of angiogenesis (*MYH9, NR4A1, SLIT3, ADAMTS12*) (14), and glucose transporters (*SLC2A3*). Complement C7 expression was increased, which has been previously reported in diabetic glomeruli (6). Changes in complement pathway genes were also observed in the cluster of cells defined by *CFH, CLDN1, VCAM1*, and *AKAP12*, which showed decreased expression of the complement inhibitory gene, *CFH* (LFC=−0.32, p=4.9e-7).

A total of 1179 endothelial cells were identified with 82 differentially expressed transcripts (Figure 2C) enriched for genes involved in vasculature development, and blood vessel endothelial cell migration. The genes included extracellular matrix components (*COL4A1*), glucose transporters (*SLC2A3*) and regulators of angiogenesis (*VEGFC, VCAM, NR4A1, MYH9, ITGB1, PRCP, TMEM204, HDAC9, MEF2C*) (15). The presence of an angiogenic signature in endothelial cells is consistent with prior reports in experimental diabetes. (16) Aberrant glomerular angiogenesis is a characteristic of diabetic nephropathy associated with glomerular hypertrophy and increased expression of endothelial VEGF (15). *VEGFC* (LFC=1.04, p=3.5e-14), and *SEMA3D* (LFC=0.55, p=6.2e-16), which belongs to a class of genes called semaphorins that direct endothelial cell motility and vascular patterning (17) were both increased. SEMA3D signals via PI3K/Akt and we detected increased expression of the phosphoinositide 3-kinase *PIK3C2B* (LFC=0.25, p=0.0009). Furthermore, diabetic endothelial cells showed increased *RAPGEF5* (LFC=0.55 p=6.2e-19), and its downstream effectors *TCF4* (LFC=0.69, p=6.7e-16) and *HDAC9* (LFC=0.62, p=0.0002), suggestive of enhanced β-catenin signaling.

### Altered Signaling Networks in the Diabetic Glomerulus

To explore alterations in intercellular signaling, differentially-expressed ligand-receptor pairs were examined in glomerular cell types (Figure 2D), which represent a subset of all interactions (Figure 2E and 2F). Diabetic mesangial cells showed increased expression of *CCN1* (LFC=0.88, p=0.014) and *SLIT3* (logFC=0.50, p=0.004). *CCN1* is a growth factor-inducible gene that regulates tissue repair via its interaction with extracellular proteins expressed by podocytes (*ITGAV, ITGB3, ITGB5*) and endothelial cells (*ITGB3*) (18). *SLIT3* modulates cell migration by interacting with *ROBO2* (19), which is also expressed by podocytes and endothelial cells. Mesangial cells expressed ligands encoded by *NAMPT, FGF1*, and *NRG3*. NAMPT regulates insulin secretion in pancreatic β-cells (20) and diabetic podocytes showed decreased expression of *INSR* (LFC=−0.31, p=0.01). NRG3 regulates cell survival pathways through its interaction with ERBB4, which was downregulated in diabetic endothelial cells (LFC=−0.86, p=0.014). Similarly, podocytes expressed *NRG3* and *EFEMP1*, which is an extracellular glycoprotein that activates the EGFR receptor (21). Diabetic endothelial cells also expressed increased *LTBP1* (LFC=0.33, p=9.3e-06), which regulates targeting of latent TGF-beta complexes.

### Infiltrating Immune Cells in Early Diabetic Nephropathy

A total of 347 leukocytes were identified and consisted of 49% T-cells, 21% B-cells, 23% mononuclear cells, and 7% plasma cells (Figure 1C). Diabetics had an approximate 7 to 8-fold increase in leukocytes compared to controls, however, this was not statistically significant (p=0.31). B-cell (n=72/74, 97%) and plasma cell (n=24/24, 100%) infiltration was unique to diabetic samples, whereas a small number of T-cells (n=150/169, 89%) and monocytes (n=62/82, 75%) were present in controls. T-cells expressed *PTPRC* (CD45), *CD247* (T-cell receptor zeta), *IL7R* (CD127), and *CD96* (Figure 1D) in addition to markers of activation and maturation (*CD28, ZAP70, THEMIS*) and the pro-survival gene, *BCL2*. Mononuclear cells expressed *FCGR2A* (CD32), *IL2RA* (CD25), *CSF1R, CSF2R*, and *TNFRSF1B*, which is a biomarker for diabetic nephropathy (22). Overall, mononuclear cells showed enrichment for genes downstream of interferon gamma signaling, including *IFNGR1, TNFRSF1B, HLA-DRB1, HLA-DRB5*, and *HLA-DQB1* in addition to the T-cell co-stimulatory molecule, *CD86*. B-cells expressed *PTPRC* (CD45), *MS4A1* (CD20), *PAX5*, and to a lesser extent, *CD19*. Plasma cells expressed *CD38* and *SDC1* (CD138).

Due to the small number of leukocytes present in controls, diabetic samples were compared to an integrated reference composed of 2 publicly available peripheral blood mononuclear cell (PBMC) datasets (23, 24). The integrated dataset enhanced our ability to identify NK cell, T-cell and monocyte subsets (Figure S3) while measuring changes in a panel of inflammatory markers (25). The kidney risk inflammatory signature (KRIS) is a recently described panel of 17 circulating plasma proteins that predict progression of diabetic nephropathy (25). We saw increased expression of *TNFRSF21* (LFC=1.12, p=7.6e-58) in the infiltrating diabetic CD14+ monocyte subset (Figure 3A), which was one of the few KRIS markers that showed a correlation between enhanced urinary excretion and end-stage renal disease (25). *ILR1* was increased in CD16+ monocytes and antigen presenting cells, and *IL18R1* was increased in CD4+ and CD8+ T-cells. These data suggest that infiltrating immune cells contribute to the production of KRIS markers.

**Figure 3.**
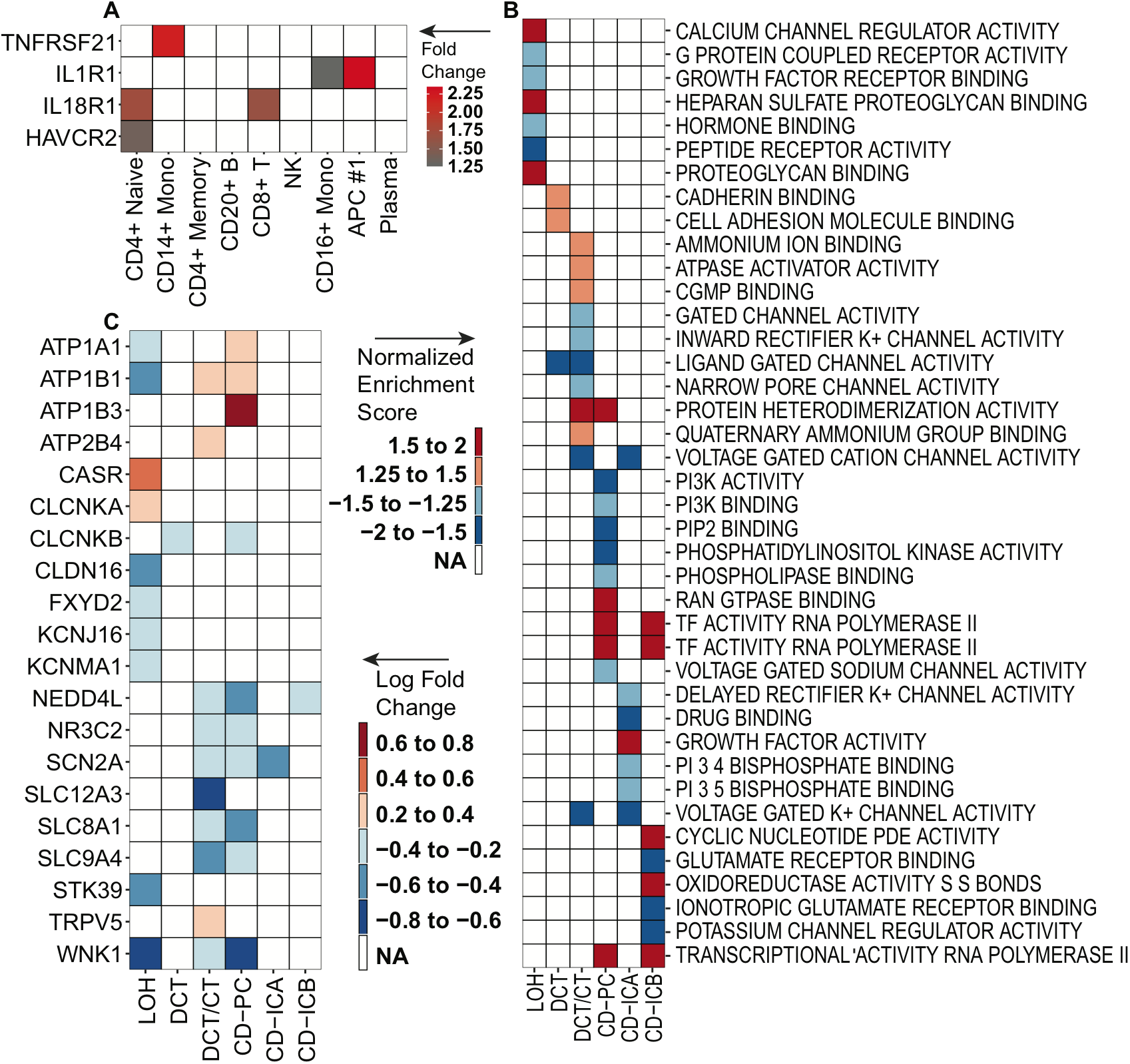
Differential expression of predictive biomarkers and ion transport pathways. – A) Leukocyte subsets extracted from the integrated dataset were interrogated for differential expression of a panel of inflammatory markers (KRIS). B) Cells from the loop of Henle (LOH) to the collecting duct underwent gene set enrichment analysis using the R package fgsea and were mapped to gene ontology terms C) Differentially expressed genes involved in ion transport in the distal nephron were identified using Seurat

### Early Gene Expression Changes in the Diabetic Proximal Convoluted Tubule and Loop of Henle

A total of 6518 proximal convoluted tubule (PCT) cells were identified with 284 differentially expressed genes, enriched for regulators of angiogenesis, cellular response to TGF-beta signaling, positive regulation of the JNK cascade, positive regulation of NF-ĸB transcription factor activity, semaphorin-plexin signaling, platelet-derived growth factor receptor signaling, and response to oxidative stress. The diabetic PCT had increased expression of the albumin transporter, *LRP2* (LFC=0.31, p=2.0=−55), and regulators of angiogenesis: *NRP1* (LFC=0.46, p=2.1e-77), *VCAM1* (LFC=0.64, p=7.6e-66), and *HIF1A* (LFC=0.45, p=2.8e-45) (26). There was a shift in metabolism characterized by up-regulation of *PCK1* (LFC=0.42, p=2.6e-28) and down-regulation of *INSR* (LFC=−0.31, P=2.0E-17). Phosphoenolpyruvate carboxykinase 1 (PCK1) is a control point for regulation of gluconeogenesis (27) and ammoniagenesis (28). Multiple intracellular signaling pathways were altered, including increased *APP* (LFC=0.34, p=6.0e-43), which binds death receptor 6 encoded by *TNFRSF21*, (29) and *NRG3* (LFC=0.38, p=2.7e-26), which binds ERBB4.

A total of 3788 cells in the loop of Henle were identified with 338 differentially expressed genes, enriched for regulation of sodium ion transmembrane transporter activity, adherens junction organization, response to glucocorticoids, regulation of canonical Wnt signaling, and angiogenesis. Na+/K+-ATPase (NKA) subunits encoded by *ATP1A1* (LFC=−0.32, p=4.5e-28), *ATP1B1* (LFC=−0.40 P=9.5E-70) and *FXYD2* (30) (LFC=−0.31, p=9.1e-41) were decreased. *WNK1* (LFC=−0.73, p=9.2e-81) and its downstream effector, *STK39* (LFC=−0.58, p=3.6e-86) were also decreased, which regulate Na+-K+-2Cl-cotransporter (NKCC2), albeit modestly (31). Nevertheless, because sodium and potassium transport in the TAL are driven by the in-series operation of NKCC2 and the NKA, the collective changes are expected to reduce Na+ and K+ reabsorption. Inhibition of transport in the TAL is expected to be compounded by the reduction in the basolateral potassium channel, *KCNJ16* (LFC=−0.29, p=3.2e-33) and an increase in the calcium sensing receptor, *CASR* (LFC=0.42, p=4.2e-51), which inhibits Na+-K+ reabsorption and drives calcium excretion by decreasing NKCC2, the ROMK channel (32) and CLDN16, which forms tight junctions by oligomerizing with CLDN19 (33). *CLDN16* (LFC=−0.58, p=2.0e-69) was decreased, which leads to increased sodium delivery to the collecting duct, increased fractional excretion of potassium, and impaired calcium and magnesium reabsorption in the thick ascending limb (34, 35).

### Diabetes induces gene expression changes that promote potassium secretion in the late distal convoluted tubule and principal cells

A total of 1652 late distal convoluted tubule (DCT2) and connecting tubule cells were present with 225 differentially expressed genes (Figure 3C), enriched for plasma membrane calcium ion transport, regulation of sodium ion transport, cellular response to hypoxia, MAPK cascade, and angiogenesis. Increased expression of the apical calcium-selective channel, *TRPV5* (LFC=0.30, p=0.002), and the basolateral plasma membrane calcium ATPase (PMCA), *ATP2B4* (LFC=0.32, p=4.5e-05), in DCT/CT likely result as a consequence of a reduction in transport within the thick ascending limb (36) as a compensatory mechanism to prevent excessive urinary calcium loss. There was also decreased *NEDD4L* (LFC=−0.26, p=2.8e-8) and increased *SGK1* (LFC=0.68, p=1.5e-27), which promote surface expression of ENaC.

A total of 2909 principal cells were present with 239 differentially expressed genes enriched for regulation of distal tubule development, regulation of potassium ion transmembrane transport, regulation of sodium ion transmembrane transport, regulation of JNK cascade, and angiogenesis. Diabetic principal cells had increased *ATP1A1* (LFC=0.26, p=6.1e-05), *ATP1B1* (LFC=0.37, p=4.3e-21), *ATP1B3* (LFC=0.61, p=5.9e-33), and the modulator *FXYD4* (LFC=0.27, p=0.0002), which increases NKA affinity for sodium and potassium (Figure 3C). Immunofluorescence studies also showed increased expression of NKA (Figure S4) although it was not statistically significant (Ratio = 1.40, p=0.28). Diabetic principal cells showed decreased *WNK1* (LFC=−0.60, p=1.2e-48). Alternative promoter usage of the *WNK1* gene creates a kidney specific WNK1 form, which lacks a kinase domain and is found mainly in the DCT, and the long-form, L-WNK1 that negatively regulates surface expression of the potassium secretory channel, *KCNJ1* (ROMK) in the principal cell (37, 38). GSEA showed decreased PI3K activity (Figure 3B) and the anticipated change in PIP2, which activates ROMK (39, 40), and decrease in WNK1 are upstream of mTOR regulation of potassium secretion (41) and are expected to enhance potassium secretion. The decrease in *NEDD4L* (LFC=−0.40, p=4.7e-34), which negatively regulates ENaC (42), would increase potassium secretion further. There was also increased expression of aquaporin-3 (LFC=0.31, p=8.2e-22), which is important for concentrating urine, and is increased in streptozotocin-induced diabetes (43). The increased expression of *AQP3* was also seen in immunofluorescence studies (Figure S5), but these results were not statistically significant (Ratio=1.70, p=0.20). Diabetic principal cells had decreased expression of the mineralocorticoid receptor (MR), *NR3C2* (LFC=−0.37, p=1.9e-45), which is suggestive of a conserved response to cellular injury that was observed in the late distal convoluted tubule.

There was decreased expression of the Na+/Ca++ exchanger (NCX), *SLC8A1* (LFC=−0.57, p=7.0e-28), which has been reported in experimental models of diabetes (44) and evidence for alteration of SLIT-ROBO signaling with an increase in *ROBO2* (LFC=0.98, p=8.4e=10) and a decrease in *SLIT2* (LFC=−0.27, p=3.7e-08). We observed increased expression of the transcription factor, *RUNX1*, in both the late distal convoluted tubule (LFC=0.66, p=3.3e-17) and principal cells (LFC=0.71, p=1e-17), which may promote renal fibrosis (45).

A total of 1874 type A intercalated cells and 693 type B intercalated cells were present in our sample. Increased *IL18* expression was seen in both type A (LFC=1.6, p<1e-300) and B (LFC=0.97, p<1e-300) intercalated cells compared to other kidney cell types, which is a candidate biomarker for diabetic nephropathy progression (46). Type A intercalated cells had 120 differentially expressed genes and had decreased expression of *IRS1* (LFC=−0.78, p=3.1e-51), which regulates insulin signaling and is associated with genetic susceptibility to type 2 diabetes in Pima Indians (47). Type B intercalated cells had 130 differentially expressed genes and were enriched for regulators of WNT signaling. Alteration of WNT signaling in type B intercalated cells was evidenced by decreased expression of positive regulators, including *GNAQ* (LFC=−0.31, p=0.0004), *TIAM1* (LFC=−0.40, p=1.4e-06), *LGR4* (LFC=−0.53, p=7.3e-07) and downstream transcription factors, *FOXO1* (LFC=−0.36, p=0.0001) and *FOXO3* (LFC=−0.47, p=0.0003). Furthermore, there was increased expression of the negative regulator of WNT signaling, *BICC1* (LFC=0.56, p=7.2e-06).

Ligand-receptor analysis of principal cells, and type A and B intercalated cells, revealed altered signaling pathways in the diabetic collecting duct (Figure S6). Diabetic principal cells showed a decrease in *FGFR2* (LFC=−0.35 p=4.0e-15), which is a receptor for FGF ligands produced by type A (FGF9) and B (FGF9) intercalated cells. There was also increased neuroligin 1 (LFC=0.33, p=7.1e-5), which encodes the ligand for neurexin 1 expressed by type A intercalated cells. Principal cells expressed EGF-family ligands, *NRG1* and *EGF*, which bind down-regulated *EGFR* (LFC=−0.28, p=0.006) and *ERBB3* (LFC=−0.30, p=7.8e-5) on type B intercalated cells. Netrin 1 expression was decreased in type A (LFC=−0.39, p=1.2e-5) and B (LFC=−0.33, p=0.007) intercalated cells, which binds UNC5C in type B intercalated cells (LFC=−0.34 p=7.4e-7). Netrin signaling may protect against ischemia-reperfusion injury in the kidney and contribute to angiogenesis in diabetic retinopathy (48, 49).

## Discussion

In this study, we report the first single nucleus RNA-sequencing dataset of human diabetic nephropathy. Diabetic patients had evidence of mild to moderate glomerulosclerosis and interstitial fibrosis, but preserved kidney function characteristic of early stage disease. By comparing changes in gene expression between control and diabetic samples, we were able to demonstrate up-regulation of pro-angiogenic genes and alterations in signaling pathways important for cell motility and actin cytoskeletal rearrangement. Furthermore, diabetes induced adaptive changes in the loop of Henle, late distal convoluted tubule, and principal cells that coordinate to promote potassium secretion.

The glomerulus is among the first areas in the kidney where diabetic injury becomes apparent. Adaptive changes lead to increased mesangial proliferation, matrix deposition, and an increase in glomerular basement membrane thickness. We detected increased expression of *COL4A1* and *COL4A2* in the diabetic mesangium, which may contribute to mesangial matrix expansion. These changes were accompanied by upregulation of pro-angiogenic genes like *CCN1* and the glucose transporter, *SLC2A3. CCN1*, also known as cysteine-rich angiogenic inducer (CYR61), is a secreted protein that promotes endothelial cell adhesion by interacting with extracellular matrix proteins expressed by podocytes and endothelial cells (18). Abnormal angiogenesis contributes to the progression of diabetic nephropathy and is characterized by upregulation of pro-angiogenic factors (15). We observed increased expression of *VEGFC* in the diabetic endothelium, which has been implicated in podocyte survival (50). Diabetic podocytes and proximal tubule cells also showed evidence of exposure to hyperglycemia, characterized by down-regulation of the insulin receptor.

A number of pathways were upregulated in multiple cell types, including WNT/β-catenin, MAPK/JNK, and EGF signaling. Alteration of WNT/β-catenin signaling was seen in the endothelium, loop of Henle, and type B intercalated cells. Similarly, alteration of MAPK/JNK signaling was seen in the proximal convoluted tubule, late distal convoluted tubule and connecting tubule, and principal cells. JNK signaling has been associated with insulin resistance and renal fibrosis (51). Diabetic endothelial cells showed decreased expression of *ERBB4* and diabetic principal cells showed decreased expression of *EGFR* and *ERBB3*. Decreased urinary EGF is a biomarker for diabetic nephropathy progression (52). The majority of *EGF* expression was detected in the loop of Henle and distal collecting tubule and although we did not detect a change between diabetic and control samples, our data suggest that principal cells may represent one of the targets.

Ligand-receptor analysis revealed multiple pathways that may be conserved between different cell types distributed throughout the kidney. Podocytes showed increased expression of *NTNG1* and *UNC5D*, which is a signaling pathway that directs axon growth and neuronal development while promoting cell survival (53). *NTNG1* expression has been previously reported in podocytes in the developing kidney, but its role in diabetic nephropathy is unknown. We also observed decreased expression of a related ligand, *NTN1*, in both type A and B intercalated cells, suggesting location-dependent variations in netrin signaling.

Early changes in gene expression in the loop of Henle, late distal convoluted tubule, and principal cells collectively promote potassium secretion (Figure 4) while decreasing paracellular calcium and magnesium reabsorption (Figure 5). The loop of Henle showed decreased expression of the Na+/K+-ATPase (NKA) subunits, *ATP1A1, ATP1B1*, and *FXYD2*, and the basolateral potassium channel, *KCNJ16*, in addition to decreased expression of *WNK1*, and its downstream effector, *STK39* (SPAK), which regulate activity of the apical Na+-K+-2Cl-cotransporter (NKCC2). Decreased NKA, KCNJ16 and NKCC2 activity in the loop of Henle are expected to impair transcellular sodium and potassium reabsorption and decrease paracellular reabsorption of calcium and magnesium. This would be exacerbated by the observed increased expression of the calcium sensing receptor (*CASR*) and decreased expression of *CLDN16*, which regulates tight junction permeability. These changes were accompanied by increased expression of *SGK1* and decreased expression of an important regulator of ENaC and potassium secretion, *NEDD4L* in the collecting duct. Interestingly, patients with type 1 or type 2 diabetes and microalbuminuria show significantly decreased activity of NKA in erythrocytes (54, 55) due to glucose-dependent inactivation (56), which is similar to findings in ob/ob mice (57). Reduced NKA activity may result from loss of early compensatory mechanisms and explain the well-known propensity of patients with diabetic nephropathy to develop hyperkalemia and type 4 renal tubular acidosis. Alternatively, this potassium handling response may reflect nephron loss with consequent requirement for higher potassium secretion per remaining nephron. In contrast, the late distal convolute tubule and principal cells showed increased expression of NKA, which promotes potassium secretion via ROMK, which is expected to be upregulated in response to decreased PI3K (41), increased PIP2 and decreased WNK1 signaling (58). In our study, the net effect of the changes seen in the loop of Henle, late distal convoluted tubule, and principal cells are expected to promote potassium secretion and may represent an adaptive response to early diabetic kidney injury.

**Figure 4.**
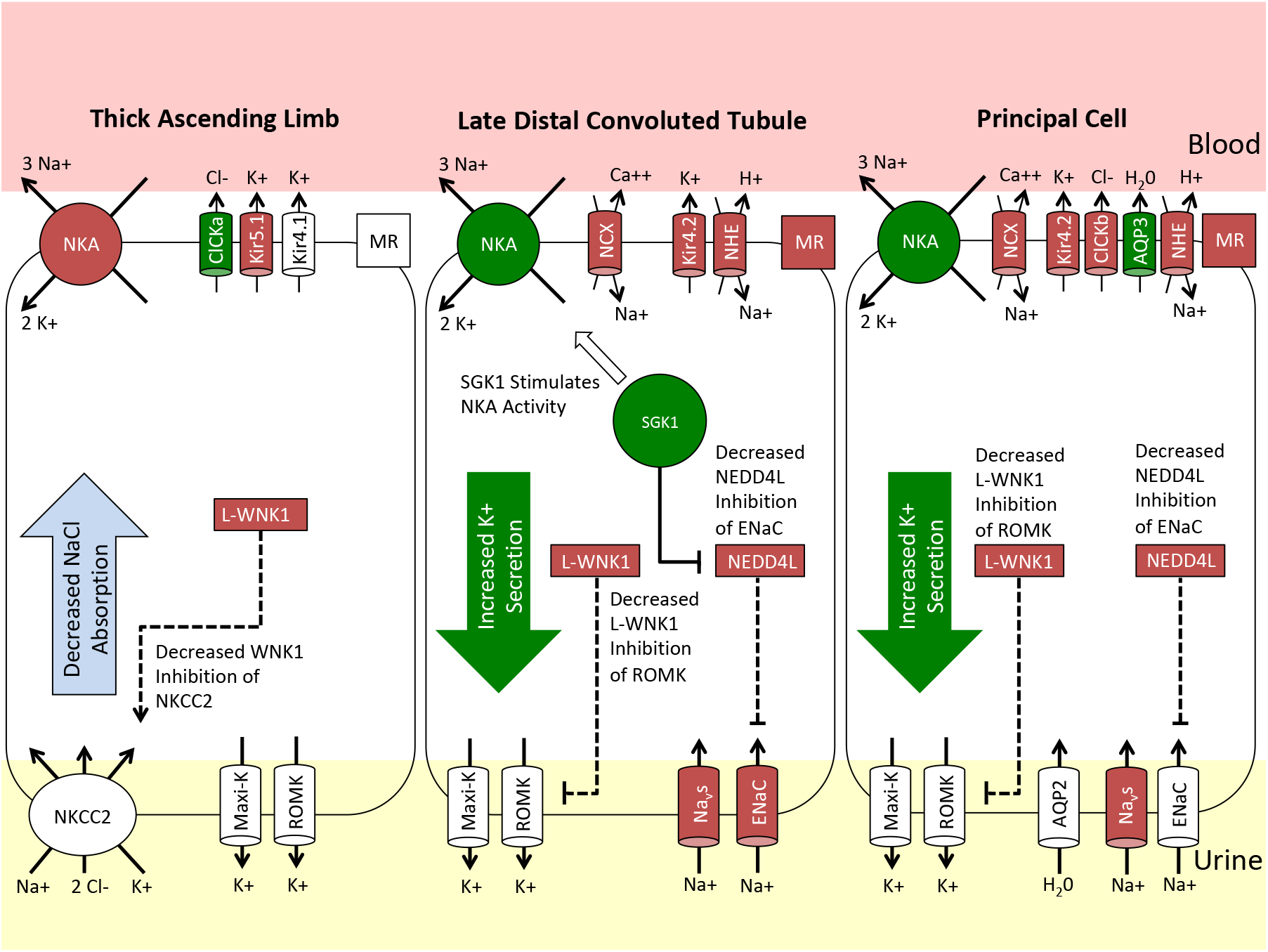
Coordination of the distal nephron to promote potassium secretion. Genes are depicted as upregulated (green fill), downregulated (red fill) or no significant change (white fill) relative to control. The loop of Henle showed decreased expression of the Na+/K+-ATPase (NKA) subunits, *ATP1A1, ATP1B1*, and *FXYD2*, and the basolateral potassium channel, *KCNJ16* (Kir5.1), in addition to decreased expression of *WNK1*, and its downstream effector, *STK39* (SPAK), which regulate activity of the apical Na+-K+-2CI- cotransporter (NKCC2). Decreased NKA, KCNJ16 and NKCC2 activity in the loop of Henle are expected to impair transcellular sodium and potassium reabsorption. These changes were accompanied by increased expression of NKA and *SGK1* with decreased expression of an important regulator of ENaC and potassium secretion, *NEDD4L*, in the collecting duct.

**Figure 5.**
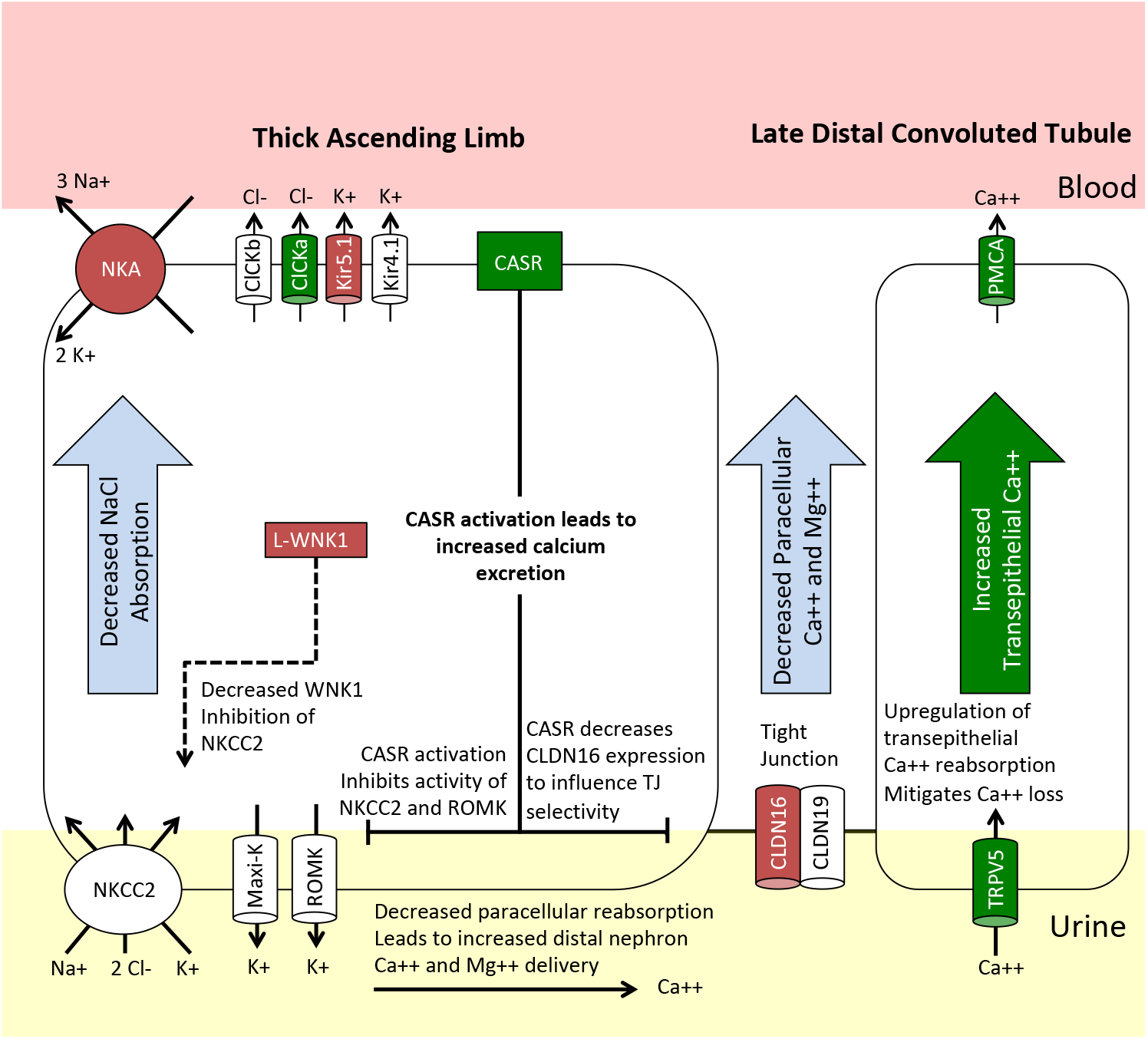
Decreased paracellular reabsorption of calcium and magnesium. - Genes are depicted as upregulated (green fill), downregulated (red fill) or no significant change (white fill) relative to controls. Decreased NKA, KCNJ16 (Kir5.1) and NKCC2 activity in the loop of Henle are expected to impair transcellular sodium and potassium reabsorption and decrease paracellular reabsorption of calcium and magnesium. This would be exacerbated by the observed increased expression of the calcium sensing receptor (*CASR*) and decreased expression of *CLDN16*, which regulates tight junction permeability. Increased expression of the apical calcium-selective channel, *TRPV5*, and the basolateral plasma membrane calcium ATPase, (PMCA), in the late distal convolutd tubule likely result as a compensatory mechanism to prevent excessive urinary calcium loss.

Immune cell infiltration is a well-known feature of diabetic nephropathy. (59) We observed an increased number of T-cells, B-cells, plasma cells, and mononuclear cells in two of three diabetic samples. The two patients with an increased number of immune cells also had mild to moderate proteinuria and interstitial fibrosis. Infiltrating mononuclear cells expressed markers downstream of interferon gamma signaling, including interferon gamma receptors (*IFNGR1* and *IFNGR2*), HLA class II histocompatibility antigens (*HLA-DRB1, HLA-DRB5, HLA-DQA1*), and *TNFRSF1B*, which has been implicated as a biomarker for diabetic nephropathy (60). Comparison of our dataset to publicly available peripheral blood mononuclear datasets showed that infiltrating CD14+ monocytes have increased *TNFRSF21*, which is a urinary marker of diabetic nephropathy progression (25).

A limitation of this study is the relatively small number of patients and future studies would benefit from increasing the sample size to better delineate inter-individual variation and capture the full range of disease severity. Our results lay the foundation for such efforts.

## Materials and Methods

### Tissue Procurement

Kidney tissue was obtained from patients undergoing partial or radical nephrectomy for renal mass at Brigham and Women’s Hospital (Boston, MA) under an established IRB protocol. Non-tumor cortical tissue was frozen or retained in OCT for future studies.

### Single-Nuclei Isolation and Library Preparation

Nuclei were isolated with Nuclei EZ Lysis buffer (NUC-101; Sigma-Aldrich) supplemented with protease inhibitor (5892791001; Roche) and RNase inhibitor (N2615; Promega and AM2696; Life Technologies). Samples were cut into <2-mm pieces and homogenized using a Dounce homogenizer (885302–0002; Kimble Chase) in 2 ml of ice-cold Nuclei EZ Lysis buffer, and they were incubated on ice for 5 minutes with an additional 2 ml of lysis buffer. The homogenate was filtered through a 40-μm cell strainer (43–50040–51; pluriSelect) and then centrifuged at 500×g for 5 minutes at 4°C. The pellet was resuspended and washed with 4 ml of the buffer, and then, it was incubated on ice for 5 minutes. After another centrifugation, the pellet was resuspended in Nuclei Suspension Buffer (1× PBS, 1% BSA, and 0.1% RNase inhibitor), filtered through a 5-μm cell strainer (43–50020–50; pluriSelect), and counted. The 10× Chromium single-cell protocol provided by the manufacturer (10× Genomics) was used to generate high-quality cDNA libraries.

### Bioinformatics and Data Analysis

Single nucleus sequencing data was processed with zUMIs (v2.0). Low-quality barcodes and UMIs were filtered using the internal read filtering algorithm and mapped to the human genome (hg38) using STAR (2.6.0). We quantified unique intronic and exonic reads to generate a raw count matrix, which was imported to Seurat. Genes expressed in >3 nuclei and nuclei with at least 500 genes were retained. Seurat objects were subsequently normalized and scaled. The number of principal components was estimated using the PCElbowPlot function.

An integrated dataset was created using canonical correlation analysis and the RunMultiCCA function with highly variable genes curated from individual Seurat objects. We obtained a new dimensional reduction matrix by aligning the CCA subspaces and performed clustering. Differential gene analysis was performed on individual clusters and visualized with Seurat. Publicly available peripheral blood mononuclear cell datasets (3k PBMCs from a Health Donor Cell Ranger 1.1.0 and 4k PBMCs from a Health Donor, Cell Ranger 2.1.0) were downloaded from the 10x Genomics website and integrated with the leukocyte subset to analyze KRIS markers (25).

### Ligand-receptor Interaction Analysis

To study ligand-receptor interactions, we used a draft network published by Ramilowski *et al* (61). We examined glomerular or tubulointerstitial cell types and required that i) the ligand, receptor, or both were differentially expressed and ii) its cognate pair was expressed in the partner cell type.

### Functional Gene Set Enrichment Analysis

Gene set enrichment analysis was performed with the R package fgsea with default parameters. Genes were ranked within clusters by multiplying avg_logFC by −log_10_(p_val) obtained from comparing diabetic and control samples with the FindMarkers function in Seurat. The human GO molecular function database was downloaded using msigdbr.

### Immunofluorescence Studies

Formalin-fixed paraffin embedded tissue sections were deparaffinized and underwent antigen retrieval. Sections were washed with PBS, blocked with 10% normal goat serum (Vector Labs), permeabilized with 0.2% Triton-X100 in PBS and incubated overnight with primary antibodies for AQP3 (LSBio, LS-B8185), ATP1A1 (abcam, EP1845Y), AQP2 (Santa Cruz, sc-9882), or V-ATPase (Santa Cruz, sc-55544) followed by staining with secondary antibodies (FITC-, Cy3, or Cy5-conjugated, Jackson ImmunoResearch). Sections were stained with DAPI (4’,6’- diamidino-2-phenylindole) and mounted in Prolong Gold (Life Technologies). Images were obtained by confocal microscopy (Nikon C2+ Eclipse; Nikon, Melville, NY). AQP3 and ATP1A1 were quantified in ImageJ by using the color threshold and measure functions to calculate integrated fluorescence density. Total corrected target immunofluorescence was calculated by subtracting the total principal cell area multiplied by the background fluorescence from the integrated fluorescence density. Results were compared between control and diabetic samples using a two-tailed students t-test.

## Acknowledgements

This work was supported by NIDDK Diabetic Complications Consortium (www.diacomp.org) grants DK076169 (to B.D.H.), DK115255 (to B.D.H.), 173970 from the Chan Zuckerberg Initiative (to B.D.H), DK104308 (to S.S.W. and B.D.H.), DK054231 (to P.A.W.) and the Leducq Fondation (to P.A.W.).

