## Supplemental Information for "The Single Cell Transcriptomic Landscape of Early Human Diabetic Nephropathy"

**This PDF file includes:**

Figs. S1 to S6
Tables S1 to S2

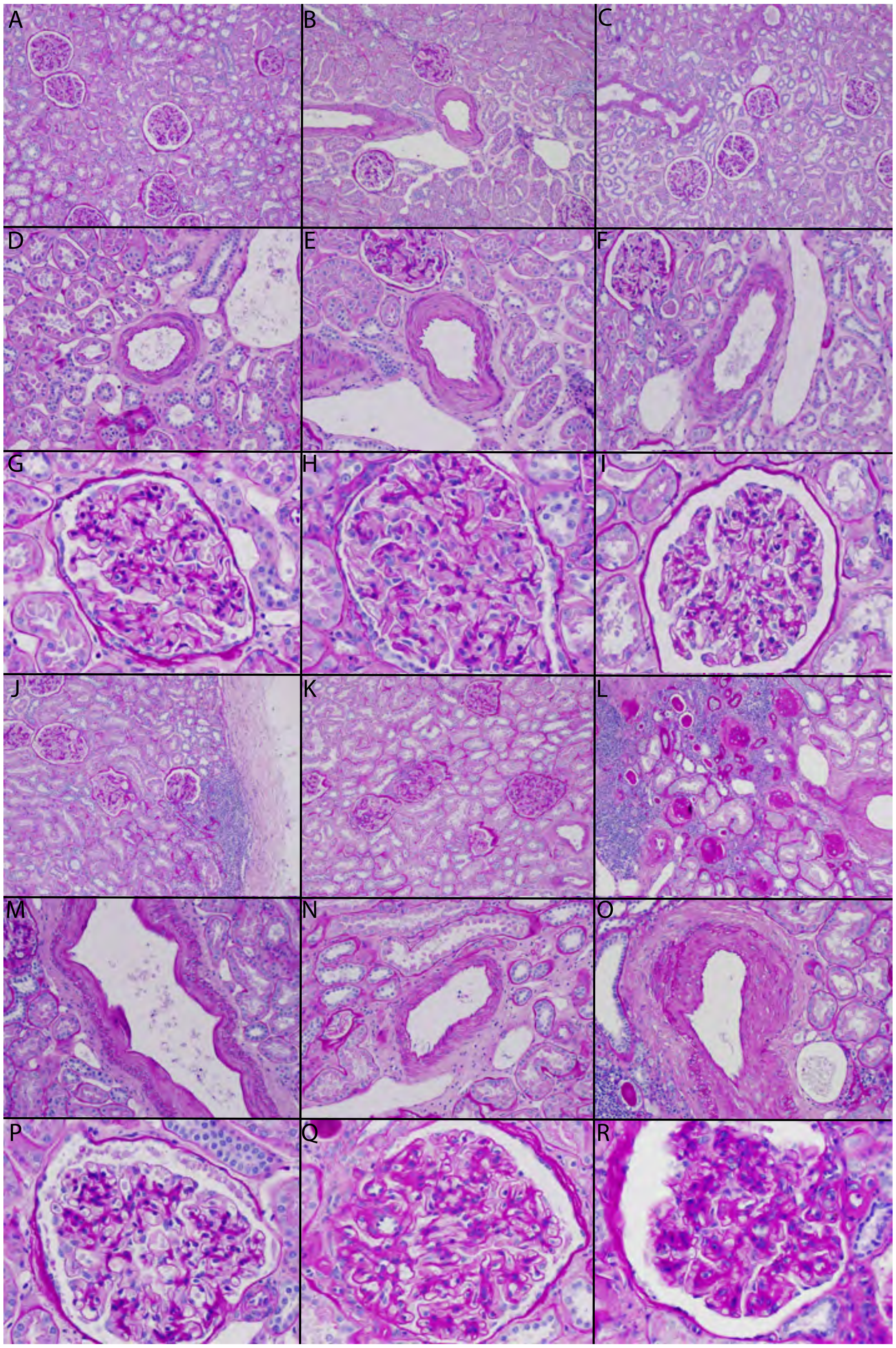


**Fig. S1.** Representative H&E images of control (A-I) and diabetic (J-R) samples. Low-power images (100x) of control samples (A-C) show no evidence of glomerulosclerosis, interstitial fibrosis, or immune cell infiltrate. Low-power images of diabetic samples (J-L) show patchy glomerulosclerosis, interstitial fibrosis, and immune cell infiltrate. Medium-power images (200x) of control (D-F) and diabetic (M-O) vessels show mild to moderate intimal sclerosis. High-power images (400x) of control glomeruli (G-I) appear normal, whereas diabetic glomeruli (P-R) show mesangial expansion and glomerular basement membrane thickening.

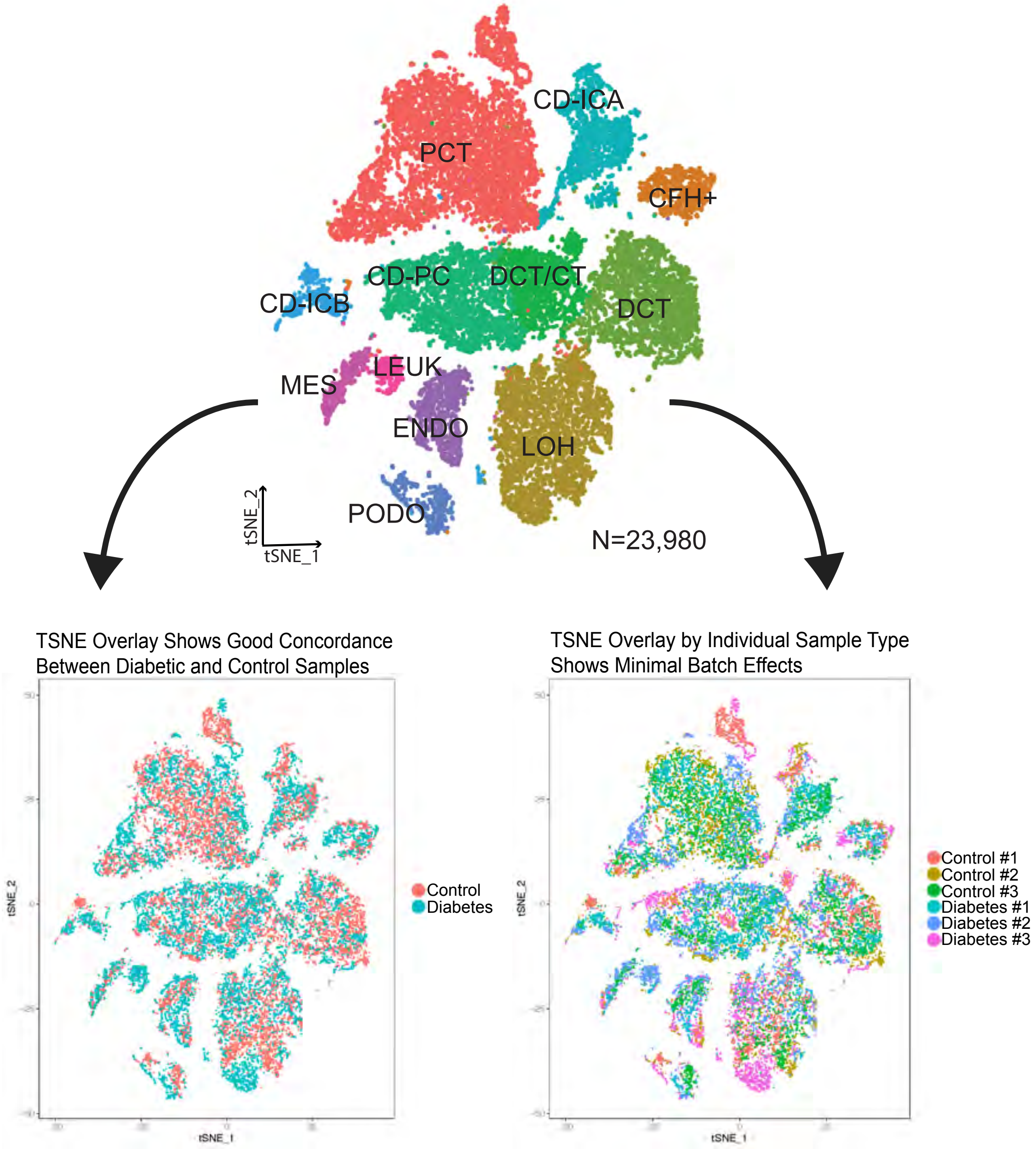


**Fig. S2.** TSNE plots of integrated dataset separated by sample type (Control vs. Diabetes) and sample of origin (Control #1-3 and Diabetes #1-3).

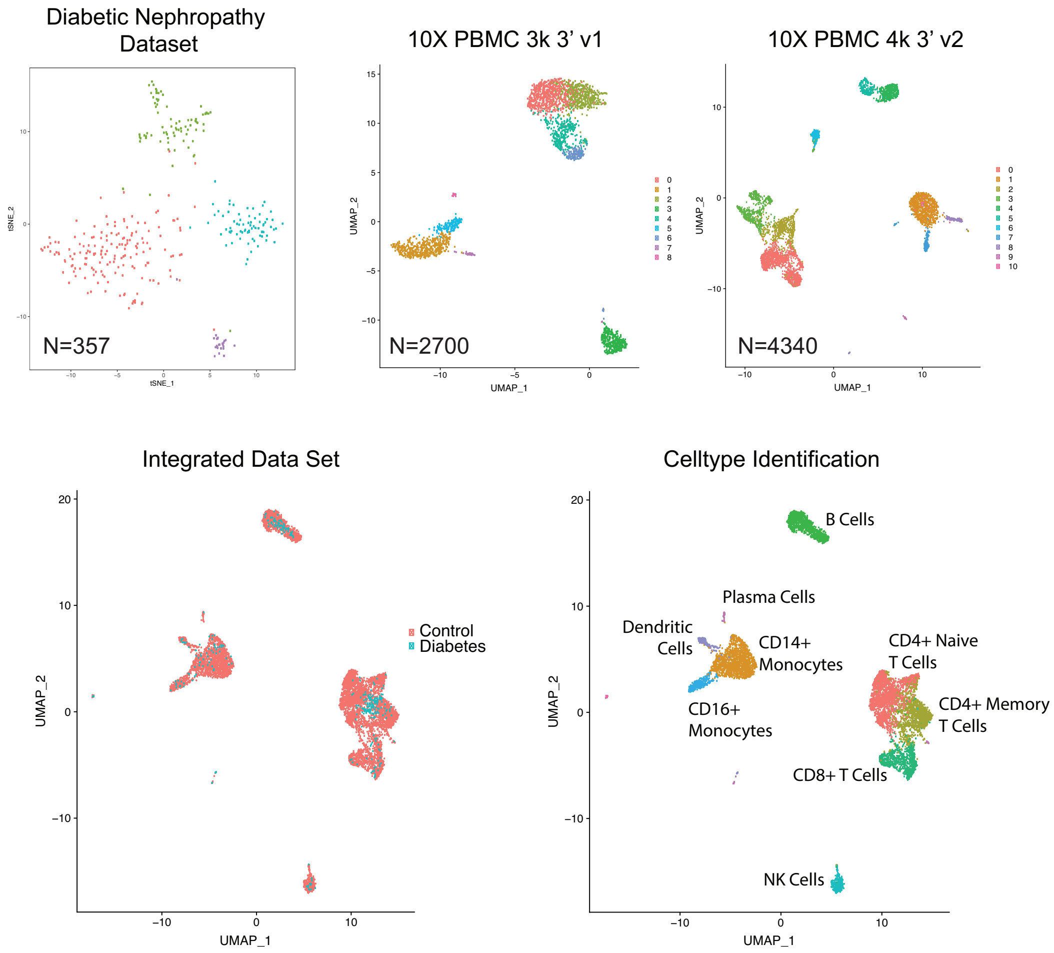


**Fig. S3. Comparison of leukocyte subset obtained from patients with diabetic nephropathy to 2 publicly-available datasets downloaded from 10x genomics**. Leukocytes from the diabetic nephropathy dataset were extracted into a separate Seurat object using the SubsetData function and integrated with pbmc datasets (3k PBMCs from a Health Donor Cell Ranger 1.1.0 and 4k PBMCs from a Health Donor, Cell Ranger 2.1.0) using Seurat 3.0. Differential gene expression was performed within leukocyte subsets.

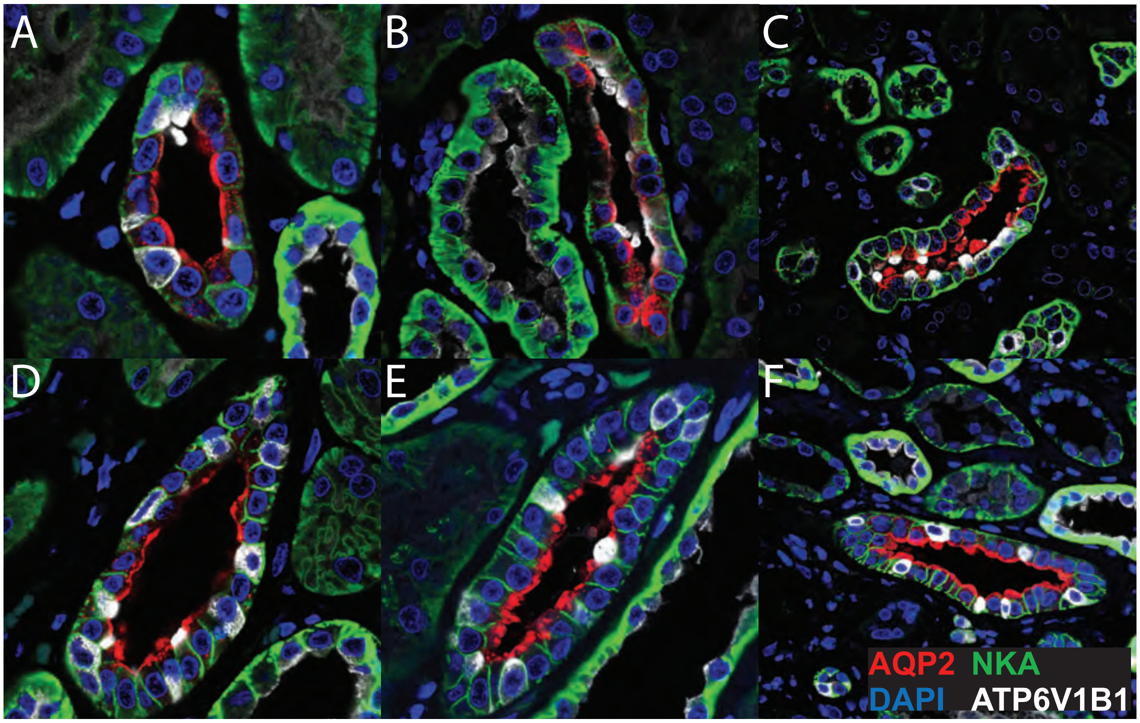


Fig. S4. **Representative images of expression of Na+/K+ ATPase (NKA) in control (A-C) and diabetic (D-F) principal cells.** Formalin-fixed paraffin embedded sections were deparaffinized and stained for AQP2 (red), Na+/K+ ATPase subunit ATP1A1 (green) and ATP6V1B1 (white) following antigen retrieval and imaged on a confocal microscope. Na+/K+ ATPase expression was quantified in AQP2+ principal cells using ImageJ. Each image represents an individual patient.

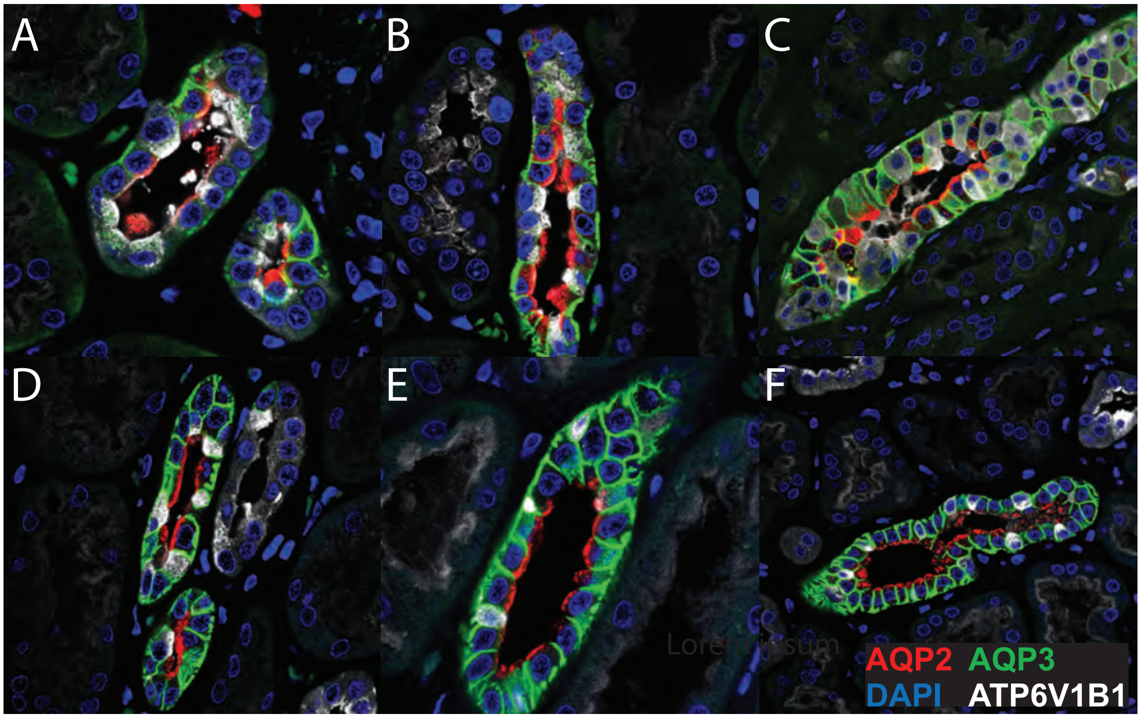


Fig. S5. **Representative images of expression of AQP3 in control (A-C) and diabetic (D-F) principal cells.** Formalin-fixed paraffin embedded sections were deparaffinized and stained for AQP2 (red), AQP3 (green) and ATP6V1B1 (white) following antigen retrieval and imaged on a confocal microscope. AQP3 expression was quantified in AQP2+ principal cells using ImageJ. Each image represents an individual patient.

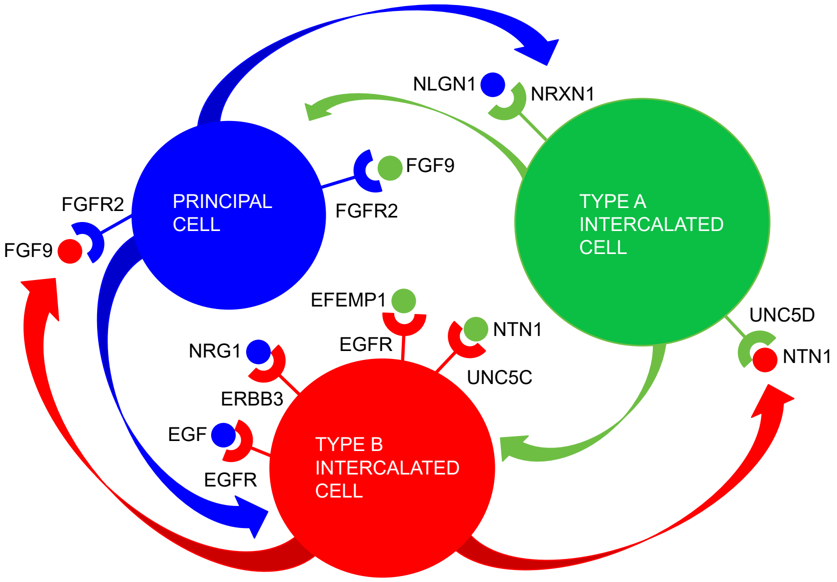


Fig. S6. **Differentially expressed ligand-receptor interactions in the collecting duct.** To study ligand-receptor interactions, we used a draft network published by Ramilowski *et al* (1). We examined collecting duct cell types and required that i) the ligand, receptor, or both were differentially expressed and ii) its cognate pair was expressed in the partner cell type. Differential expression was assessed using the FindMarkers function in Seurat with default settings.

**Table S1. Demographic, laboratory and pathology data for patient samples**

| | Control Patients (n=3) | | | Diabetic Patients (n=3) | | |
|---|---|---|---|---|---|---|
| Age, Gender | 54M | 62M | 61F | 74M | 52M | 57F |
| A1c | NA | NA | NA | 6.7 | 7.3 | 9.7 |
| sCr | 1.28 | 1.21 | 0.89 | 1.26 | 1.03 | 0.7 |
| Proteinuria | NA | NA | NA | Yes | 2+ | No |
| Glomerulosclerosis | None (<10%) | None (<10%) | None (<10%) | Moderate (26-50%%) | Mild (11-25%) | None (<10%) |
| IFTA | 1-10% | 1-10% | 1-10% | 11-25% | 11-25% | 1-10% |
| Arteriosclerosis | Moderate | Moderate | Mild | Mild | Moderate | Mild |

**Table S2. Number of nuclei and number of genes and UMI per nucleus for each sample**

| | Control 1 | Control 2 | Control 3 | Diabetes 1 | Diabetes 2 | Diabetes 3 | Total |
|---|---|---|---|---|---|---|---|
| Nuclei Accepted | 3740 | 3917 | 6203 | 3771 | 3806 | 2543 | 23,980 |
| Mean No. Genes per Nucleus | 4046 | 1354 | 2114 | 3142 | 1902 | 2689 | 2541 |
| Mean No. UMI per Nucleus | 12736 | 2734 | 4717 | 9806 | 4813 | 6561 | 6894 |
